# Interneuron functional diversity in the mouse accessory olfactory bulb

**DOI:** 10.1101/552463

**Authors:** Marina A. Maksimova, Hillary L. Cansler, Kelsey E. Zuk, Jennifer M. Torres, Dylan J. Roberts, Julian P. Meeks

## Abstract

In the mouse accessory olfactory bulb (AOB), inhibitory interneurons play an essential role in gating behaviors elicited by sensory exposure to social odors. Several morphological classes have been described, but the full complement of interneurons remains incomplete. In order to develop a more comprehensive view of interneuron function in the AOB, we performed targeted patch clamp recordings from partially-overlapping subsets of genetically-labeled and morphologically-defined interneuron types. *Gad2* (GAD65), *Calb2* (calretinin), and *Cort* (cortistatin)-cre mouse lines were used to drive selective expression of tdTomato in AOB interneurons. *Gad2* and *Calb2-*labeled interneurons were found in the internal, external, and glomerular layers, whereas *Cort*-labeled interneurons were enriched within the lateral olfactory tract (LOT) and external cellular layer (ECL). We found that external granule cells (EGCs) from all genetically-labeled subpopulations possessed intrinsic functional differences that allowed them to be readily distinguished from internal granule cells (IGCs). EGCs showed stronger voltage-gated Na^+^ and non-inactivating voltage-gated K^+^ currents, decreased I_H_ currents, and robust excitatory synaptic input. These specific intrinsic properties did not correspond to any genetically-labeled type, suggesting that transcriptional heterogeneity among EGCs and IGCs is not correlated with expression of these particular marker genes. Intrinsic heterogeneity was also seen among AOB juxtaglomerular cells (JGCs), with a major subset of *Calb2*-labeled JGCs exhibiting spontaneous and depolarization-evoked plateau potentials. These data identify specific physiological features of AOB interneurons types that will assist in future studies of AOB function.

**Significance Statement:** The mouse accessory olfactory bulb (AOB) plays a critical role in processing social chemosensory information. Several morphologically-identified types of AOB inhibitory interneurons are thought to refine and restrict information flow from the AOB to its downstream targets in the limbic system. However, little is known about the electrophysiological and transcriptional diversity among AOB interneuron types. We systematically investigated intrinsic electrophysiological diversity across 5 AOB cell populations in three transgenic mouse lines. Analysis of 26 intrinsic physiological features revealed feature combinations associated with identified morphological AOB cell types, but few associated with the transgenic lines we studied. The results provide quantitative information about functional diversity in AOB interneurons and provide an improved foundation for future studies of AOB circuit function.

## Introduction

The mammalian brain utilizes a diverse complement of interneuron types to achieve its overall function. In well-studied neural circuits like the hippocampus, neocortex, and retina, interneuron types have been painstakingly identified and catalogued over a period of decades using a combination of morphological and electrophysiological metrics (Freund and Buzsaki, 1996; Somogyi and Klausberger, 2005; Burton, 2017). Transgenic and viral technologies allow selective labeling and manipulation of specific interneuron types in olfactory circuits, facilitating research into functional subdivisions among genetically-defined types (Parrish-Aungst et al., 2007; Eyre et al., 2009; Huang et al., 2013; Kato et al., 2013; Miyamichi et al., 2013; Tatti et al., 2014; Burton et al., 2017). Importantly, many neural circuits, including the AOB, lack a comprehensive “taxome” that links genetic, morphological, and functional properties of interneurons. A more complete and quantitative description of the interneuron classes in the AOB would provide the ground knowledge useful for building a deeper understanding of AOB circuit function.

The mouse accessory olfactory bulb (AOB), the first neural circuit in the mammalian accessory olfactory system (AOS, also called the vomeronasal system), is a critical regulator of mouse behavior. Although analogous cell types appear to be present in the main olfactory bulb (MOB), it is clear that the AOB and MOB have distinct organizations (Jia et al., 1999; Taniguchi and Kaba, 2001; Araneda and Firestein, 2006; Castro et al., 2007; Shpak et al., 2012; Smith et al., 2015; Gorin et al., 2016). In the quest to achieve a better understanding of AOB function and its impact on social behavior, establishing a taxome of AOB interneuron is an important step.

A comprehensive morphological survey of the AOB neuronal types identified up to 6 distinct interneuron types in the AOB (Larriva-Sahd, 2008). Importantly, among the identified cell types in the AOB is a large set of spiny interneurons called external granule cells (EGCs) that have not been studied in depth. EGCs reside in the external cellular layer (ECL) alongside mitral cells (MCs), the only projection neurons of the AOB. EGCs, like JGCs and IGCs, appear to lack axons, and are thought to form reciprocal dendro-dendritic synapses with MCs. EGCs have elaborate radial dendritic arbors studded with long-necked gemmules or spines, and as such likely make extensive lateral connections in the circuit. However, physiological data concerning the intrinsic and synaptic features of these cells are needed to build informed hypotheses about their function.

Here, we present electrophysiological data recorded from 150 genetically-and morphologically-defined AOB interneurons across all of the major AOB sublaminae, including JGCs, EGCs, and IGCs. In order to avoid explicit bias in our choice of interneurons from which to record, and to test for potential links between marker gene expression and function, we made targeted recordings from largely non-overlapping sets of genetically-labeled AOB interneurons (via *Calb2-cre*, *Gad2-cre*, and *Cort-cre* transgenic mice). We utilized a battery of intrinsic electrophysiological assays and multidimensional analyses to identify functionally similar groups. We found that IGCs, EGCs, and JGCs possess characteristic physiological features that are largely invariant across genetically-labeled groups. EGCs, in particular, possess a combination of physiological features that indicate a potential role in broad MC inhibition. These data show clear functional subdivisions across morphologically-defined AOB interneuron classes, but also reveal that physiological features of these interneurons are shared across genetically distinct types.

## Materials and Methods

### Mice

75 adult male and female mice aged 6-12 weeks were utilized in this study. All animals were housed in the care of the [Author University] Animal Resource Center, and were given food and water *ad libitum* before euthanasia. All studies were performed in accordance with the [Author University] animal care committee’s regulations. Animals were housed with a 12-12 light-dark cycle, and slices were taken at Zeitgeber time of 20-24 hours, corresponding to the latter part of time in darkness for the mice.

To generate offspring expressing the fluorescent protein tdTomato in genetically-restricted types, we crossed *Gad2^tm2(cre)Zjh^/J* (Jackson Labs Stock 010802, “*Gad2-cre”*), *B6(Cg)-Calb2^tm1(cre)Zjh^/J*, (Jackson Labs Stock 010774, “*Calb2-cre”*), or *Cort^tm1(cre)Zjh^/J* (Jackson Labs Stock 010910, “*Cort-cre”*) animals to *B6;129S6-Gt(ROSA)26Sor^tm9(CAG-tdTomato)Hze^/J* (Jackson Labs Stock 007905, *ROSA26-loxP-STOP-loxP-tdTomato* or “Ai9”) animals (Madisen et al., 2010; Taniguchi et al., 2011). Genotyping was performed by Transnetyx, Inc. All transgenic animals were heterozygous for both knock-in transgenes. All animals were backcrossed for multiple generations with C57Bl/6J animals prior to arrival at our facility.

### Immunohistochemistry

Animals were anesthetized lightly with isofluorane, then injected with ketamine/xylazine cocktail (120mg/kg ketamine, 16mg/kg xylazine) to induce deep anesthesia. Animals were transcardially perfused with cold phosphate buffered saline (PBS) followed by 4% paraformaldehyde in PBS (PFA-PBS). Brains were then removed from the skull, bisected at the midline, and post-fixed in PFA-PBS for 2-12 h. Hemispheres were then cryoprotected in PBS with 25% w/v sucrose for 12-24 h. Hemispheres were aligned with the sagittal plane, then frozen in a block of OCT compound (TissueTek) for 1-2 h, then sectioned on a cryostat (Leica CM3050 S) at 20 µm.

Sections were rinsed 3x with PBS in 24-well plates, then permeabilized by incubating for 2 h in PBS containing 0.3% Triton-X 100. Sections were blocked in PBS containing 0.1% Triton-X 100 containing 10% goat serum (“primary block”) for 2-4 h. Primary antibodies were diluted into primary block, and sections were incubated overnight at 4°C, followed by a 4x rinse in PBS. Secondary antibodies were diluted into PBS containing 0.1% Triton-X 100 and 5% goat serum (“secondary block”). Sections were exposed to secondary antibodies for 2 h, followed by a final 4x PBS rinse. Negative controls omitted primary antibodies (not shown). Sections were mounted on slides, coverslipped using Fluoromount-G (SouthernBiotech) and sealed with nail polish.

Images of immunostained sections were taken at 40x (1.3 NA) on a Zeiss LSM 510 confocal microscope. All images for a given staining batch were imaged with matched laser power and photomultiplier tube gain. For display purposes, maximum projection images of 5-10 frames (1 µm per frame) were created with ImageJ, then stitched manually in Adobe Photoshop. Quantification of cell density was performed using ImageJ.

### Acute slice preparation

Standard artificial cerebrospinal fluid (aCSF) contained (in mM): NaCl 125, KCl 2.5, CaCl_2_ 2, MgCl_2_ 1, NaHCO_3_ 25, NaH_2_PO_4_ 1.25, myo-inositol 3, Na-pyruvate 2, Na-ascorbate 0.4, glucose 25. Animals were anesthetized lightly with isofluorane and then rapidly decapitated into ice-cold aCSF with an additional 9 mM MgCl_2_ (10 mM MgCl_2_ total). Brains were extracted in cold slicing buffer, and the anterior portion of the brain containing the olfactory bulbs and frontal neocortex was isolated from the rest of the brain. This anterior brain tissue was separated into two hemispheres and then embedded in aCSF containing 4% agarose in aCSF. The tissue was then affixed to an angled slicing tray with tissue glue in a vibrating microtome (Leica). Slices were cut at an angle of at approximately 12 degrees off-sagittal such that the caudal/medial aspect of the olfactory bulbs was cut first. We find this angle to better preserve the long IGC neurites. After vibrosectioning, slices were immediately placed into a holding chamber containing standard aCSF containing 0.5 mM kynurenic acid. Slices were allowed to recover for at least 30 minutes prior to placing in the recording chamber.

### Electrophysiology

Electrophysiological experiments were performed using a custom epifluorescence upright microscope (Nikon). Brain slices were placed in a large volume tissue chamber (Warner Instruments) warmed to 33°C by a temperature controller (Warner Instruments). tdTomato-expressing neurons were identified by imaging with a 40x objective (Olympus) and digital camera (Point Grey) with dim fluorescent illumination (X-Cite 200DC). Patch electrodes were pulled on a horizontal Flaming/Brown-style puller (P1000, Sutter Instruments) using thin-walled borosilicate glass (TW150, World Precision Instruments). Pipette resistances ranged from 6 to 10 MΩ, and optimal recordings were achieved with 8-9 MΩ electrodes. Internal solution for intrinsic characterization experiments contained (in mM): K-gluconate 115, KCl 20, HEPES 10, EGTA 2, MgATP 2, Na_2_GTP 0.3, Na-phosphocreatine 10, with a pH of 7.37. Internal solution for voltage clamp experiments on mitral cells contained (in mM): Cs-methanesulfonate 130, NaCl 4, CaCl_2_ 0.5, EGTA 5, HEPES 10, MgATP 4, Na_2_GTP 0.3, QX314 5, with a pH of 7.35.

Electrophysiological signals were amplified using a MultiClamp 700B amplifier controlled via pClamp 10 software (Molecular Devices/Axon Instruments). Signals were acquired at 20 kHz via a Digidata 1440 analog/digital converter (Molecular Devices/Axon Instruments). Signals were filtered at 10 kHz at acquisition until processing with custom software written in MATLAB (see Data Analysis subheading). Access resistance was monitored at the beginning and end of each experiment, and recordings during which the access resistance increased by more than 50% or exceeded 40 MΩ were excluded from further analysis. Series resistance was not compensated.

For interneuron morphological reconstructions, Alexa Fluor 488 hydrazide (100-200 µM, ThermoFisher/Molecular Probes) was included in the recording solution. In these experiments, slices were mounted on a Acerra two-photon microscope (ThorLabs) and filled for 20-30 minutes. Morphologies were reconstructed from 100-300 µm optical stacks (1-2 µm/optical section) after performing manual morphological tracing using the Simple Neurite Tracer plugin in ImageJ (Longair et al., 2011).

### Data analysis

To assess intrinsic electrophysiological features, we subjected patched AOB neurons to a series of current clamp and voltage clamp challenges. Immediately after achieving the whole cell configuration, each cell’s resting membrane potential (V_rest_) was measured in current clamp mode. To standardize measurements across cells with different V_rest_, we injected steady-state currents to maintain each cell’s membrane potential (V_m_) between −0 and −0 mV. Based on initial measurements of input resistance (R_input_), we empirically determined the amplitude of hyperpolarizing current that adjusted V_m_ by approximately −0 mV (to −0 to −0 mV). After determining this initial current injection amplitude, we generated a cell-specific 10-sweep Clampex protocol that applied increasingly depolarizing 0.5 s square current pulses, starting with the initial injection amplitude. For example, if the initial current injection was determined to be −0 pA, the 10-sweep protocol would have current injection increments of +20 pA (*i.e.*, −0 pA, −0 pA, −0 pA,…,+80 pA). If the initial depolarization was determined to be - 125 pA, the protocol would include increments of +25 pA, etc. This strategy allowed us to objectively challenge cells with widely varying V_rest_ and R_input_. In voltage clamp, cells were initially held at −0 mV, and a series of 12 voltage command steps (0.5 s in duration) were applied that spanned −0 mV to +10 mV.

For each cell, both current clamp and voltage clamp protocols were applied up to 4 times, and all reported quantities represent the mean responses across repeated trials. Twenty-six specific intrinsic parameters were extracted from each cell using custom software written in MATLAB. A description of these parameters in Fig. 7A and the formulas used to calculate them is presented in Table 1. For Cluster analysis using a custom algorithm based on the mean-shift strategy (Comaniciu and Meer, 2002) was performed on normalized values. Values were normalized by the 95^th^ percentile absolute value for each physiological feature across the entire cell population (N=150 cells). After normalization, resulting values were truncated to the range of −0 to 1.

**Table 1.** Parameters used for multidimensional analysis.

| Shorthand | Description | Mode | Method | Reference |
| --- | --- | --- | --- | --- |
| Vrest | resting membrane potential | I Clamp | direct measurement | N/A |
| Cm | membrane capacitance | I Clamp | see reference | Golowasch, et al, 2009 |
| Rm | input resistance |  |  |  |
| Ncomps | number of model compartments |  |  |  |
| Ih_sag | hyperpolarization-induced depolarizing potential | I Clamp | $I_{H,sag} = V_{init} - V_{ss}$ | N/A |
| Ih_current | hyperpolarization-induced depolarizing current | V Clamp | $I_{H,curr} = I_{init} - I_{ss}$ | N/A |
| EPSC_freq | mean spontaneous EPSC frequency | V Clamp | direct measurement after waveform sorting | Hendrickson, et al, 2008 |
| EPSC_amp | mean spontaneous EPSC amplitude | V Clamp |  |  |
| EPSC_tau | mean spontaneous EPSC decay constant | V Clamp | least-squares exponential decay | N/A |
| S_freq_1 | Spiking frequency - Lv. 1 | I Clamp | direct measurement | N/A |
| S_freq_2 | Spiking frequency - Lv. 2 | I Clamp | direct measurement | N/A |
| S_freq_3 | Spiking frequency - Lv. 3 | I Clamp | direct measurement | N/A |
| S_freq_4 | Spiking frequency - Lv. 4 | I Clamp | direct measurement | N/A |
| S1_slope | Initial action potential rising slope | I Clamp | initial spike derivative peak | Meeks, et al, 2005 |
| S1_thresh | Initial action potential threshold | I Clamp | Vm @ 10% of initial spike derivative |  |
| S_accom_1 | Spike rate accomodation - Lv. 1 | I Clamp | $\frac{S_{init} - S_{final}}{S_{init} + S_{final}}$ | N/A |
| S_accom_2 | Spike rate accomodation - Lv. 2 | I Clamp |  |  |
| S_accom_3 | Spike rate accomodation - Lv. 3 | I Clamp |  |  |
| S_accom_4 | Spike rate accomodation - Lv. 4 | I Clamp |  |  |
| Na_curr_1 | Voltage-gated sodium current amplitude - Lv. 1 | V Clamp | direct measurement | N/A |
| Na_curr_2 | Voltage-gated sodium current amplitude - Lv. 2 | V Clamp | direct measurement | N/A |
| Na_curr_3 | Voltage-gated sodium current amplitude - Lv. 3 | V Clamp | direct measurement | N/A |
| Na_curr_4 | Voltage-gated sodium current amplitude - Lv. 4 | V Clamp | direct measurement | N/A |
| Na_curr_5 | Voltage-gated sodium current amplitude - Lv. 5 | V Clamp | direct measurement | N/A |
| K_curr_max | Voltage-gated potassium current - maximum | V Clamp | direct measurement | N/A |
| K_curr_diff | Non-inactivating voltage-gated potassium current | V Clamp | direct measurement | N/A |

EPSCs were automatically detected and later separated from noise using a custom computer assisted waveform-based event sorting program written in MATLAB (Hendrickson et al., 2008). EPSC decay was measured by calculating the best fit single exponential for the decay period of the EPSC. Initial action potential rising slope was calculated by measuring the peak of the first derivative of voltage with respect to time (dV/dt). Threshold was defined as V_m_ at the time the dV/dt voltage reached 10% of its peak value. Membrane capacitance and input resistance were calculated according to current clamp-based multi-compartmental algorithms (Golowasch et al., 2009). Briefly, the voltage response of each cell to a hyperpolarizing current step was fit with a series of multi-exponential curves, and the best fit determined by identifying the solution with the lowest value of the Bayesian Information Criterion to avoid over-fitting.

## Results

### Transcriptional diversity among AOB interneuron populations

Several electrophysiological studies have shown that the AOB is organized quite differently than the adjacent – and superficially similar – MOB (Araneda and Firestein, 2006; Smith et al., 2009; Leszkowicz et al., 2012; Shpak et al., 2012; Taniguchi et al., 2012). Interneuron types in the MOB have been identified based on the selective expression of marker genes (Parrish-Aungst et al., 2007; Kato et al., 2013; Miyamichi et al., 2013). In the AOB, interneuron classifications have been made based on antibody staining (Jacobowitz and Winsky, 1991; Takami et al., 1992; Porteros et al., 1995). With the goal of studying the functional diversity among morphologically-and genetically-labeled AOB interneuron types, we screened through several transgenic mouse lines in which AOB interneurons could be labeled via cre-mediated genetic recombination (Fig. 1A-C). We identified three promising lines in which strong cre-mediated gene expression was observed in AOB interneurons: *Gad2-cre*, *Calb2-cre*, and *Cort-cre* (Taniguchi et al., 2011). In each line, cre expression was introduced via knock-in to the 3’ untranslated region of the gene, leaving the endogenous coding regions intact. *Cre* transgenic mice were mated to *Rosa26-loxP-STOP-loxP-tdTomato* reporter mice (” et al., 2010) to produce transgenic mice in which specific populations of AOB interneurons were fluorescently labeled (referred to as *Gad2*-tdTomato, *Calb2*-tdTomato, and *Cort*-tdTomato animals; Fig. 1A-C).

**Figure 1.**
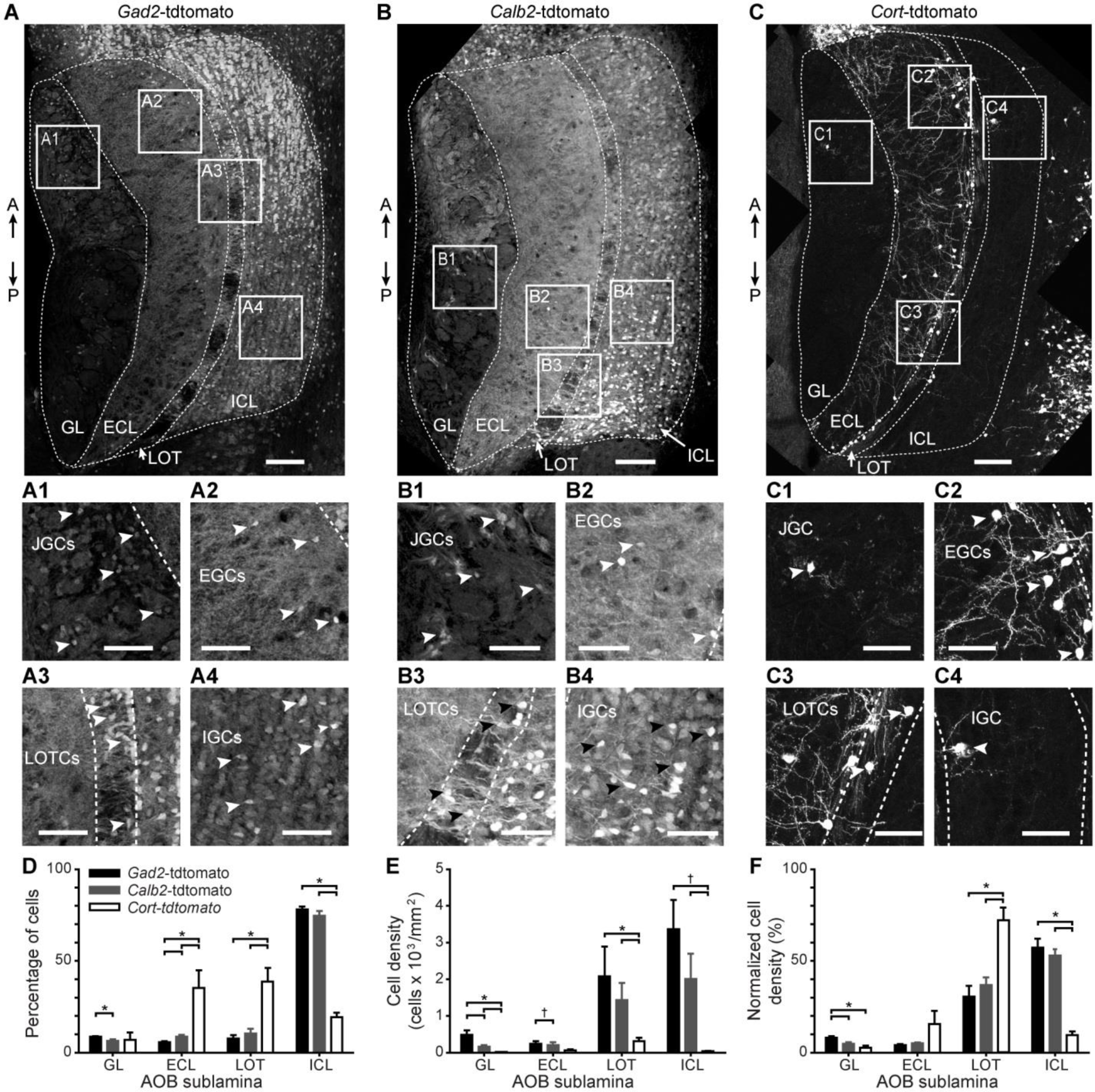
Cre-mediated genetic labeling of interneurons in all 4 AOB sublaminae. (**A**) Fluorescence micrograph of an adult male mouse AOB from a *Gad2*-tdTomato double transgenic mouse (*Gad2-*IRES-cre^+/-^ x Rosa26-loxP-STOP-loxP-tdTomato^+/-^). Boxes labeled **A1-A4** correspond to the magnified inset panels below. A↑: anterior. P↓: posterior. (**A1-A4**) Magnified views of boxes from **A** centered on the GL, highlighting JGCs (**A1**), ECL, highlighting EGCs (**A2**), LOT, highlighting LOTCs (**A3**), and ICL, highlighting IGCs (**A4**). Arrowheads indicate the locations of selected tdTomato-positive interneuron cell bodies. (**B**) Fluorescence micrograph of an adult male *Calb2*-tdTomato double transgenic mouse. (**B1-B4**) Magnified views of boxes indicated in **B**. (**C**) Fluorescence micrograph of an adult male *Cort*-tdTomato double transgenic mouse. (**C1-C4**) Magnified views of boxes indicated in **C**. (**D**) Quantification of the percentage of all tdTomato-positive cells in the GL, ECL, LOT, and ICL from *Gad2*-tdTomato (black bars), *Calb2*-tdTomato (gray bars), and *Cort*-tdTomato mice (white bars). (**E**) Quantification of the cell density of tdTomato-positive cells by layer and genotype. (**F**) Quantification of the normalized cell density of tdTomato-positive cells by layer and genotype (normalization was within genotype). GL: glomerular layer. ECL: external cellular layer. LOT: lateral olfactory tract. ICL: internal cellular layer. JGC: juxtaglomerular cell. EGC: external granule cell. LOTC: lateral olfactory tract cell. IGC: internal granule cell. Scale bars in **A-C**: 100 µm. Scale bars in magnified panels: 50 µm. * indicates p < 0.05 and † indicates 0.05 < p < 0.1, (Student’s unpaired, two-tailed *t*-test).

We observed strong tdTomato expression in all AOB cell layers in adult *Gad2*-tdTomato and *Calb2*-tdTomato mice (Fig. 1A-B), consistent with antibody staining observations (Jacobowitz and Winsky, 1991; Takami et al., 1992; Parrish-Aungst et al., 2007). The percentage of both *Gad2*-tdTomato-and *Calb2*-tdTomato-positive cells was greatest in the internal cellular layer (ICL) of the AOB, which contains neurons called internal granule cells (IGCs), the most plentiful neuronal type in the AOB (N = 4; Fig. 1D). The overall density of *Gad2*-tdTomato+ and *Calb2*-tdTomato+ neurons was greatest in the ICL and adjacent LOT (N = 4; Fig. 1E). *Gad2*-tdTomato+ and *Calb2*-tdTomato+ populations, despite densely labeling IGCs, appeared to label partially non-overlapping populations. The pattern of *Gad2*-tdTomato+ and *Calb2*-tdTomato+ IGCs varied along the anterior-posterior axis of the AOB, with *Gad2*-tdTomato+ IGCs enriched in the anterior AOB (aAOB) and *Calb2*-tdTomato+ IGCs enriched in the posterior AOB (pAOB; Fig. 1A and 1B). These anterior-posterior biases were most prominent in the lateral-most regions of the AOB (N = 3).

In contrast to the broad *Gad2*-tdTomato and *Calb2*-tdTomato labeling, we observed highly selective labeling of external granule cells (EGCs), a recently-described AOB interneuron type, in *Cort*-tdTomato double transgenic mice (Larriva-Sahd, 2008)(Fig. 1C). EGCs are multi-polar cells that possess elaborate, spine-laden dendrites and, like IGCs, lack an apparent axon (Larriva-Sahd, 2008). The percentage of *Cort*-tdTomato+ interneurons was greatest in the ECL and adjacent LOT, but these neurons were scarce in the glomerular layer (GL,) and ICL (N = 4, Fig. 1D). The overall density of *Cort*-tdTomato labeling was much lower than *Gad2*-or *Calb2*-tdTomato populations (N = 4, Fig. 1E). The percentage of *Cort*-tdTomato+ cells, when normalized by laminar area (normalized cell density, Fig. 1F), revealed that these cells were tightly clustered along the border of the ECL and LOT (N = 4). These three transgenic driver lines thus label interneurons in all AOB sublaminae, and suggested transcriptional diversity even within morphological types.

### Electrophysiological characteristics of AOB IGCs

IGCs are the predominant neuronal type in the AOB, and have been studied more thoroughly than any other interneuron in this circuit (Jia et al., 1999; Taniguchi and Kaba, 2001; Araneda and Firestein, 2006; Smith et al., 2009; Taniguchi et al., 2012; Zimnik et al., 2013; Smith et al., 2015; Hu et al., 2016). These interneurons are thought to play prominent roles in AOB function, including plasticity (Bruce, 1959; Kaba and Keverne, 1988; Brennan et al., 1990; Okutani et al., 2002; Araneda and Firestein, 2006). Despite the likely importance of AOB IGCs for AOB computations, their electrophysiological features have not been extensively studied. Given the transcriptional diversity within the IGC population (Fig. 1), we hypothesized that IGCs, as a class, comprise multiple, functionally-distinct subpopulations. We further hypothesized that type distinctions may be identified by thoroughly evaluating IGC intrinsic physiological features, a strategy that has been successful in other neural circuits (Cadwell et al., 2017; Tavakoli et al., 2018; Ting et al., 2018).

We developed an unbiased assay involving a series of electrophysiological challenges in whole cell current clamp and voltage clamp, and applied these challenges to 58 IGCs of multiple transcriptional types (Fig. 2). This approach revealed several consistent features among IGCs, most notably a propensity for spike accommodation upon moderate somatic depolarization (Fig. 2B-F) and a prominent HCN-channel mediated I_H_ “sag” potential that was blocked by the HCN channel antagonist ZD7288 (10 µM)(Fig. 2B-D, Fig. 2G). IGCs displayed moderate rates of spontaneous excitatory postsynaptic currents (sEPSCs) mediated by AMPA-and NMDA-type ionotropic glutamate receptors and metabotropic glutamate receptors (Fig. 2K-N). Despite these relatively consistent features, we observed remarkable intrinsic feature variability among IGCs (Fig. 2E-J), consistent with the hypothesis that the IGC class may comprise several distinct subpopulations.

**Figure 2.**
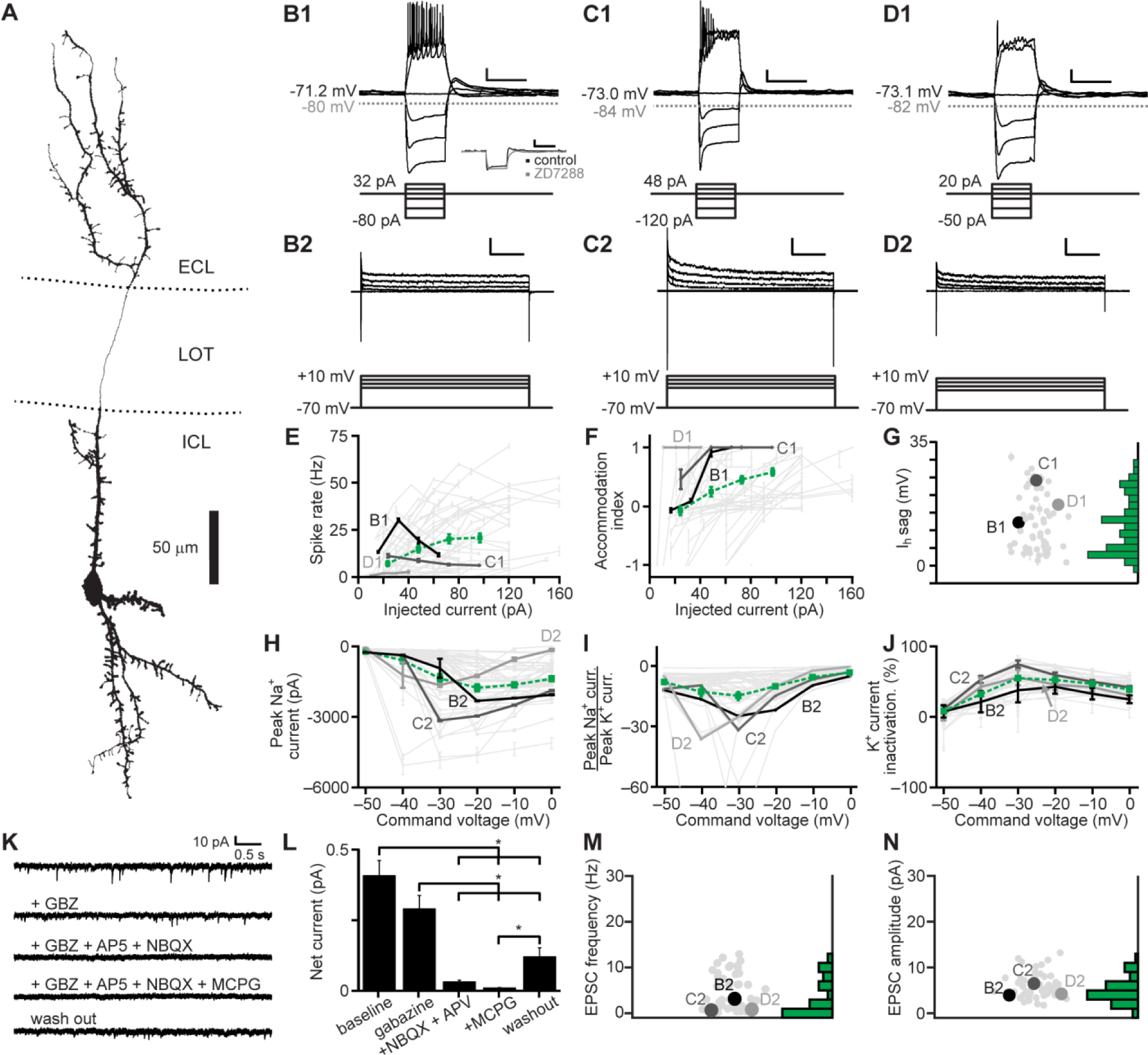
Electrophysiological properties of AOB IGCs. (**A**) Morphological reconstruction of an IGC dendritic arbor. ECL: external cellular layer. LOT: lateral olfactory tract. ICL: internal cellular layer. (**B1-D1**) Responses of three representative IGCs (top) to current clamp challenges (bottom). Scale bars: 10 mV, 500 ms. Inset in (B1) shows the blockade of the hyperpolarization-activated depolarizing I_H_ “sag” potential by ZD7288 (10 µM). (**B2-D2**) Responses of the same 3 IGCs (top) to a series of command potential steps from −0 mV in voltage clamp (bottom). Hyperpolarizing responses were also recorded, but are not shown. Scale bars: 500 pA, 100 ms. (**E**) Spike rate input-output curves for IGCs subjected to a series of step depolarizations in current clamp. Dashed green line indicates the mean ± standard error (N = 52). (**F**) Spike accommodation index for IGCs subjected to the same series of step depolarizations shown in Panel C (N = 52). (**G**) Maximal I_H_ sag potential for 52 IGCs. (**H**) Input-output curves for peak inward voltage-gated Na^+^ currents (N = 51). (**I**) Input-output curves for the ratio of peak voltage-gated Na^+^ to peak voltage-gated K^+^ currents (N = 52). (**J**) Input-output curve for normalized voltage-gated K^+^ current inactivation (N = 52). (**K**) Representative recordings of spontaneous synaptic currents (command potential −0 mV) before and during blockade of GABA_A_ receptors with gabazine (2.5 µM), AMPA and NMDA receptors (AP5 10 µM, NBQX 1 µM), and Type I/II mGluRs (MCPG 100 µM, N = 15). (**L**) Blockade of net spontaneous synaptic currents during pharmacological blockade (holding potential −0 mV). Asterisks indicate p < 0.05 by multiple comparisons of mean ranks, Kruskal-Wallis test. (**M**) Spontaneous EPSC frequency (N = 52). (**N**) Spontaneous EPSC amplitude (N = 52).

To test whether the variability in electrophysiological features might be associated with the different genetic/transcriptional labels (Fig. 1), we performed multidimensional analysis on a panel of 26 objectively measured intrinsic features (Cansler et al., 2017) (Table 1, Fig. 3). In this analysis we included recordings from 12 AOB MCs that had undergone the same battery of intrinsic physiological challenges. To account for the considerable variation in these physiological properties across cell types, data for each parameter were normalized by the 95^th^ percentile absolute value across all 150 recordings in this study, then truncated to fit within the range of –1 to 1 (see Materials and Methods). Analysis of a population of 42 IGCs and 12 MCs from which we had successfully extracted all 26 parameters revealed 3 macroscopic clusters, two of which exclusively contained IGCs, and the other of which was dominated by MCs (Fig. 3A). Sorting the cells within each cluster using principal components analysis (PCA) further revealed diversity within one of the two IGC clusters (Fig. 3A). This division was made clearer by performing multidimensional scaling of the matrix subjected to final clustering (Fig. 3B-C). The first cluster contained a large cohort of IGCs that demonstrated strong I_H_ conductances, fired high frequency bursts of action potentials, but failed to sustain high rates of firing in the face of moderate somatic depolarization (Fig. 3A). The second cluster was more distributed (visible in the multidimensional scaling analysis in Fig. 3B-C). The first subgroup within the second IGC cluster (Fig. 3A) contained IGCs that were better capable of sustaining high frequency spiking without strong accommodation and relatively low I_H_ conductances. The other subgroup within Cluster 2 contained cells that were poorly able to spike even with moderate to strong somatic depolarization, but possessed moderate I_H_ conductances and showed clear evidence of spontaneous glutamatergic input. The final cluster was dominated by mitral cells, which were clearly distinguishable from IGCs through large differences in several properties (Fig. 3A-C). The likely reason that the sub-clusters within Cluster 2 were not automatically identified by this clustering algorithm, which identifies cluster boundaries based on multidimensional statistical variance (Comaniciu and Meer, 2002; Turaga and Holy, 2012), is that this dataset had relatively large variance and low total number of cells.

**Figure 3.**
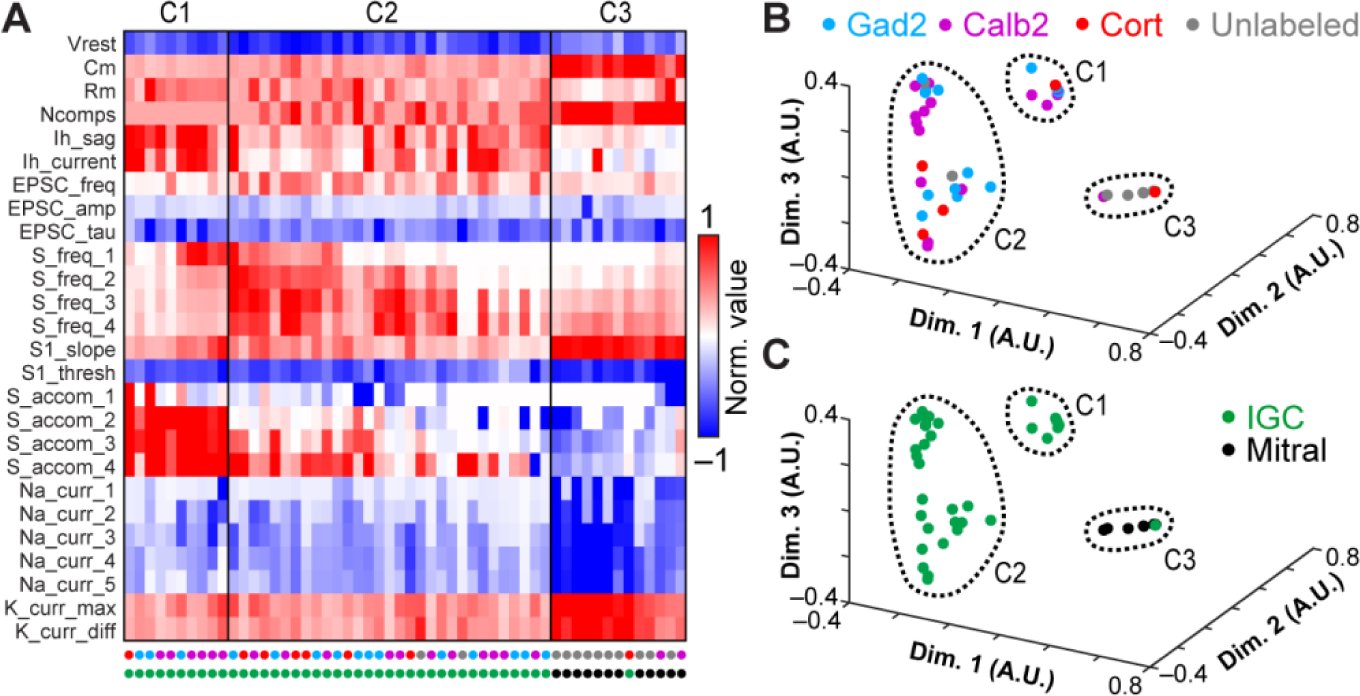
Multidimensional analysis of IGC physiological properties. (**A**) Multidimensional analysis of 26 intrinsic physiological properties was used to investigate potential relationships between different genetically-defined populations of IGCs, with mitral cells included for comparison. Each column represents the physiological profile of a single cell. Data were normalized to the 95^th^ percentile absolute value observed for each feature across 150 AOB neurons, including IGCs, EGCs, JGCs, and mitral cells. 42/58 recorded IGCs and 12/17 mitral cells contained information for all 26 parameters and were subjected to cluster analysis. Row labels refer to the intrinsic properties listed in Table 1 and column labels are color coded based on genetically-defined and morphologically-defined type. Clusters are separated by the vertical lines. (**B-C**) Nonclassical multidimensional scaling analysis of the 54 cells shown in **A**. Each symbol represents a single cell. Relative symbol positions reflect similarity across all 26 measurements (dimensions). In **B**, colored symbols indicate the genetic type of the recorded neurons; in **C** colored symbols indicate the morphological type (same color scheme as **A**). Dashed lines identify each cluster from **A**.

Having defined several subgroups among IGCs based on this multidimensional approach, we proceeded to test whether any of these clusters showed evidence of being enriched in any of the 3 transgenic lines used to label IGCs (Fig. 1). A binomial test for differential inclusion in the two IGC clusters based on genotype indicated only one such association, which was moderate enrichment of *Calb2*-tdTomato IGCs in Cluster 1 (6/10 cells, base probability 35.1%, p = 0.027). By contrast, this same analysis showed clear evidence for enrichment of mitral cells in Cluster 3 (12/13 cells, base probability 22.2%, p = 3.2 × 10^-9^). Multidimensional scaling produced no further support for the hypothesis that intrinsic physiological differences within the IGC population were strongly associated with a particular transgenic label. These results suggest that IGCs are both transcriptionally and physiologically diverse, but that these particular genetic labels do not strongly correlate with intrinsic physiological features.

### Electrophysiological characteristics of AOB EGCs

In the MOB, parvalbumin-expressing interneurons in the external plexiform layer perform MC gain scaling (Kato et al., 2013; Miyamichi et al., 2013). The AOB lacks these parvalbumin-expressing interneurons (data not shown), but possesses transcriptionally-diverse set of EGCs (*Gad2*-, *Calb2*-, and *Cort*-expressing, Fig. 1). We hypothesized that EGCs, which possess broad, densely-spine-laden dendrites (Fig. 1)(Larriva-Sahd, 2008), would possess electrophysiological properties that are distinct from IGCs, and which may vary by genetically-labeled type.

We therefore measured the intrinsic and synaptic properties of EGCs using the same protocols used for IGCs (Fig. 4). Several differences between EGCs and IGCs were apparent (Fig. 4B-D). First, resting membrane potentials for EGCs were strongly hyperpolarized (−83.6 ± 1.3 mV, N = 31) compared to IGCs (−75.6 ± 1.3 mV, N = 31, p = 1.27 × 10^-4^ Wilcoxon rank sum test). EGCs responded to depolarization with sustained 20-30 Hz spiking, showing accommodation only at the highest somatic current injections (Fig. 4E-F). EGC recordings revealed small I_H_ sag potentials that were dramatically smaller than those observed in IGCs (Fig. 4G). Consistent with the higher rates of sustained spiking in many EGCs, we observed stronger voltage gated Na^+^ and K^+^ currents in EGCs than in IGCs (Fig. 4H-I). Despite this difference in K^+^ current magnitude, EGC K^+^ currents were similar to those of IGCs in their proportion of slow inactivation (Fig. 4J). Like IGCs, spontaneous EPSCs were overwhelmingly carried by AMPA-and NMDA-type glutamate receptors (Fig. 4L), but the frequency and amplitudes of sEPSCs onto EGCs were many-fold higher than those of IGCs, indicating much denser connectivity with MCs on a per-cell basis (Fig. 4M-N).

**Figure 4.**
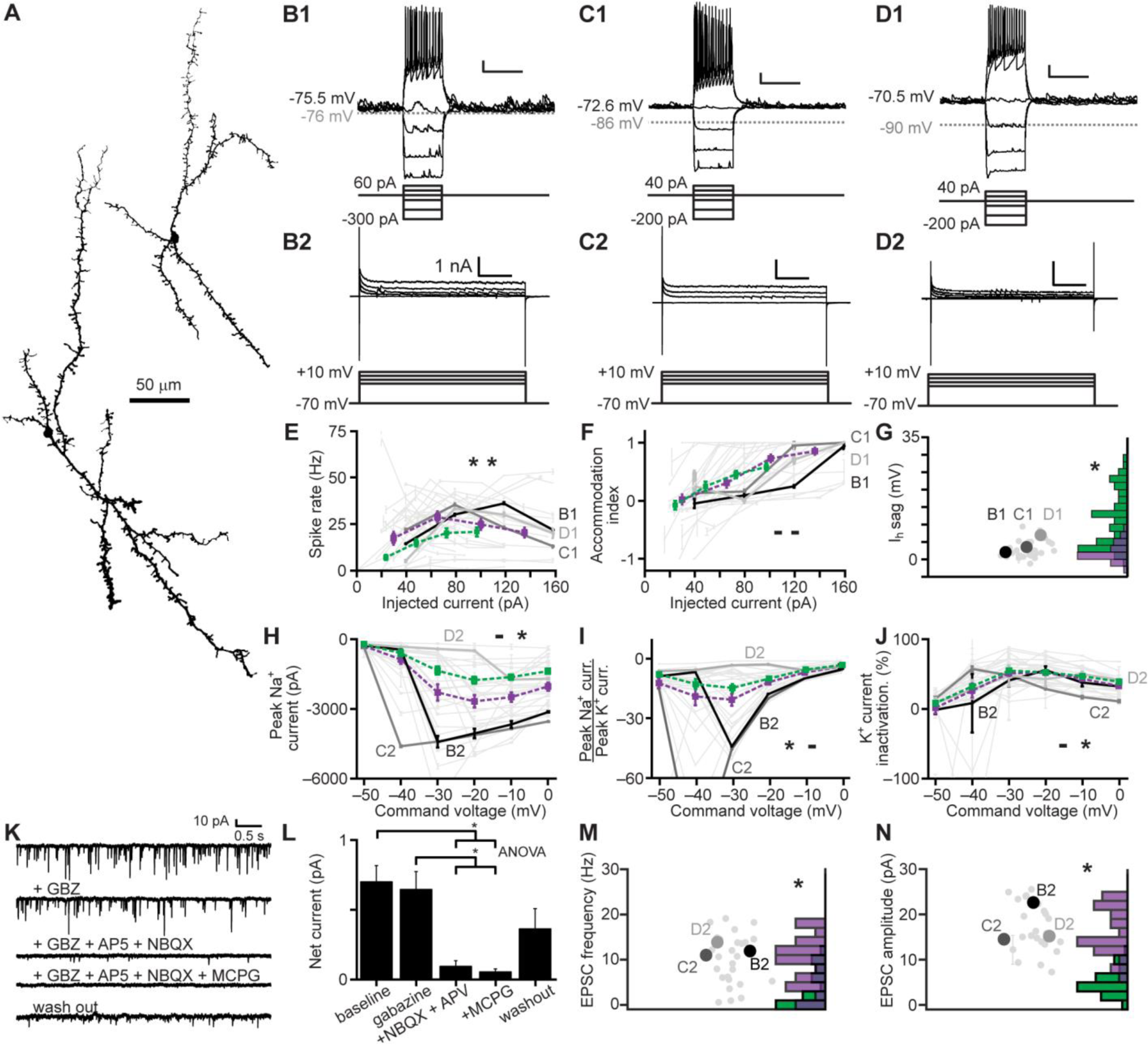
Electrophysiological properties of AOB EGCs. (**A**) EGC morphological reconstructions. (**B1-D1**) Responses of three representative EGCs to current clamp challenges. Scale bars: 10 mV, 500 ms. (**B2-D2**) Responses of the same 3 EGCs to a series of command potential steps from −0 mV in voltage clamp. Hyperpolarizing responses were also recorded, but are not shown here. Scale bars: 1 nA, 100 ms. (**E**) Spike rate input-output curves for EGCs. Dashed purple line indicates the mean ± standard error for EGCs, and dashed green for IGCs (from Fig. 2E; N for EGCs = 33, 2-way ANOVA compared to IGCs: p = 1.48 × 10^-5^ main effect of cell type, p = 0.0051 interaction between cell type and stimulus intensity). (**F**) Spike accommodation index for EGCs (N = 33, 2-way ANOVA compared to IGCs: p = 0.83 main effect of cell type. P = 0.113 interaction between cell type and stimulus intensity). (**G**) Maximal I_H_ sag potential (N = 33, compared to IGCs p = 1.48 × 10^-9^ by Wilcoxon rank sum test). (**H**) Input-output curves for peak inward voltage-gated Na^+^ currents (N = 29, 2-way ANOVA compared to IGCs: p = 0.518 main effect of cell type, p = 0.0267 interaction between cell type and stimulus intensity). (**I**) Input-output curves for the ratio of peak voltage-gated Na^+^ to peak voltage-gated K^+^ currents (N = 29, 2-way ANOVA compared to IGCs: p = 0.0048 main effect of cell type, p = 0.0916 interaction between cell type and stimulus intensity). (**J**) Input-output curve for normalized voltage-gated K^+^ current inactivation (N = 29, 2-way ANOVA compared to IGCs: p = 0.0381 main effect of cell type, p = 0.1823 interaction between cell type and stimulus intensity). (**K**) Representative recordings of spontaneous synaptic currents (command potential −0 mV) before and during application of gabazine, AP5, NBQX, and MCPG (N = 5). (**L**) Blockade of net spontaneous synaptic currents during pharmacological blockade (holding potential −0 mV). Asterisks indicate p < 0.05 by multiple comparisons of mean ranks, 1-way ANOVA. (**M**) Spontaneous EPSC frequency (N = 29, p = 2.84 × 10^-6^ Wilcoxon rank sum test) Purple bars refer to EGCs, green bars to IGCs. (**N**) Spontaneous EPSC amplitude (N = 29, p = 9.53 × 10^-13^ Wilcoxon rank sum test).

The readily-observable differences in intrinsic features suggested that EGCs might be distinguishable from IGCs when all physiological properties were considered. We performed multidimensional clustering across IGCs and EGCs by combining the physiological profiles of 29 EGCs to the group of 42 IGCs and 12 MCs analyzed in Figure 3 and re-clustering the data (Fig. 5). This analysis revealed the presence of 8 clusters within the combined dataset. Consistent with our hypothesis, Clusters 1 and 3 were overwhelmingly comprised of IGCs, Cluster 4 included fast-spiking IGCs and some EGCs, Clusters 5-7 were overwhelmingly comprised of EGCs, and Cluster 8 was dominated by MCs (Fig. 5A). As was the case with IGCs, we found little evidence for strong associations between these physiologically-derived clusters and transcriptional type (3/5 *Gad2*-tdTomato cells in Cluster 2, p = 0.022; 6/13 cells in Cluster 4, p = 0.011). On the other hand, enrichment based on morphological type was clear in all clusters except Cluster 4. These results further support the observation that EGC intrinsic physiological features are clearly distinct from IGCs, but that these features do not correlate with labeling via these 3 transgenic strategies.

**Figure 5.**
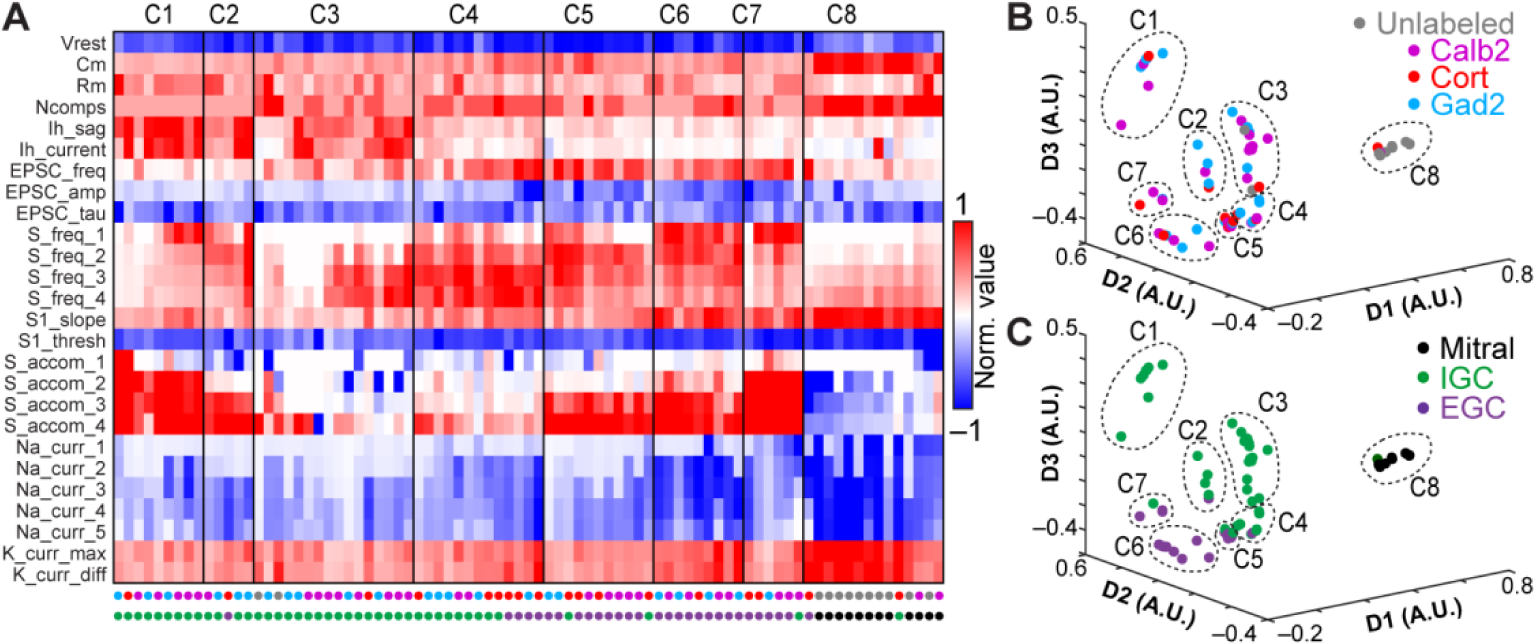
Multidimensional analysis of IGC and EGC physiological properties. (**A**) Cluster analysis of IGCs and EGCs with mitral cells included for comparison. Normalized values were calculated as per Figure 3. 29/35 recorded EGCs had information for all 26 parameters. EGC data were combined with the population of neurons in Figure 3 and re-clustered. Row labels refer to intrinsic properties listed in Table 1. Each column represents a single cell, and column labels are color coded based on genetically-defined type and morphologically-defined type. Clusters are separated by solid vertical lines. (**B-C**) Nonclassical multidimensional scaling of the 83 cells shown in **A**, colorized by genetically-defined type (**B**) and morphologically-defined type (**C**). Dashed outlines indicate approximate cluster boundaries. C1-C8 refer to the cluster definitions in **A**.

### Electrophysiological characteristics of AOB JGCs

The MOB has several interneuron types, collectively termed JGCs, which reside in and around the glomerular layer (Larriva-Sahd, 2008; Yokosuka, 2012). In the MOB, JGCs include excitatory external tufted cells, several types of inhibitory periglomerular cells, and short axon cells (Shipley and Ennis, 1996; Schoppa et al., 1998; Aungst et al., 2003; Parrish-Aungst et al., 2007; Shao et al., 2009; Liu et al., 2013; Najac et al., 2015; Burton et al., 2017; Geramita and Urban, 2017). In the AOB, very few electrophysiological inquiries have been undertaken on JGCs (Goldmakher and Moss, 2000; Hu et al., 2016). We performed electrophysiological recordings from JGCs in *Gad2*-, *Calb2*, and *Cort*-tdTomato transgenic mice, using the electrophysiological profiling strategy used for IGCs and EGCs (Fig. 6).

**Figure 6.**
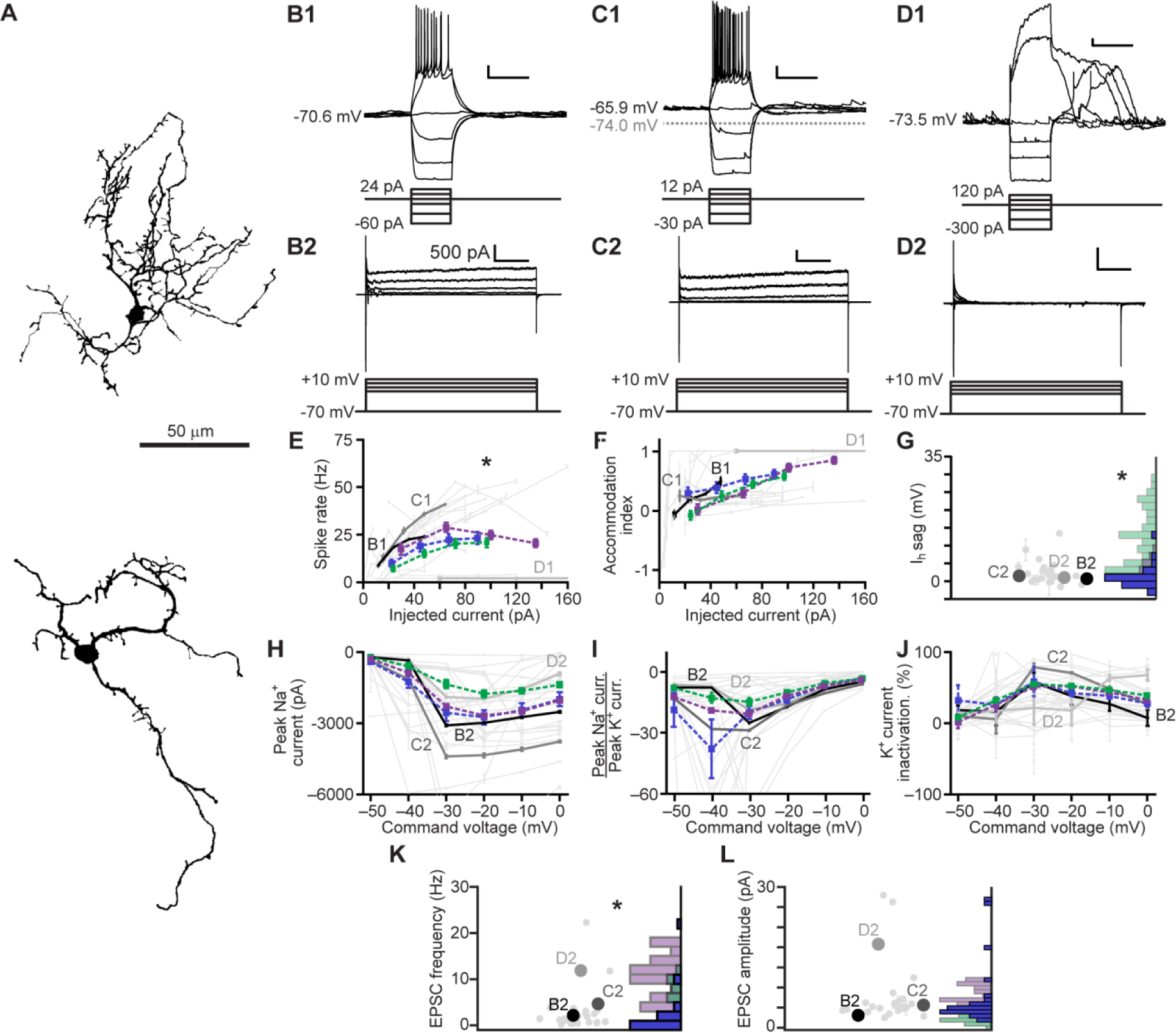
Electrophysiological properties of AOB JGCs. (**A**) JGC morphological reconstructions. (**B1-D1**) Responses of three representative JGCs (top) to current clamp challenges (bottom). Scale bars: 10 mV, 500 ms. (**B2-D2**) Responses of the same 3 JGCs (top) to a series of command potential steps from –70 mV in voltage clamp (bottom). Hyperpolarizing responses were also recorded, but are not shown here. Scale bars: 500 pA, 100 ms. (**E**) Spike rate input-output curves for JGCs. Dashed red line indicates the mean ± standard error for JGCs, dashed gray for IGCs, dashed black from EGCs (N = 27; 2-way ANOVA compared to IGCs: p = 0.414 main effect of cell type, p = 0.850 interaction between cell type and stimulus intensity; 2-way ANOVA compared to EGCs: p = 0.032 main effect of cell type, p = 0.0254 interaction between cell type and stimulus intensity). (**F**) Spike accommodation index for JGCs (N = 27, 2-way ANOVA compared to IGCs: p = 0.013 main effect of cell type. P = 0.078 interaction between cell type and stimulus intensity; 2-way ANOVA compared to EGCS: p = 0.041 main effect of cell type, p = 0.007 interaction between cell type and stimulus intensity). (**G**) Maximal I_H_ sag potential (N = 30, compared to IGCs p = 1.07 × 10^-10^, to EGCs 0.07 by Wilcoxon rank sum test). (**H**) Input-output curves for peak inward voltage-gated Na^+^ currents (N = 27, 2-way ANOVA compared to IGCs: p = 0.065 main effect of cell type, p = 0.323 interaction between cell type and stimulus intensity; 2-way ANOVA compared to EGCs: p = 0.346 main effect of cell type, 0.456 interaction between cell type and stimulus intensity). (**I**) Input-output curves for the ratio of peak voltage-gated Na^+^ to peak voltage-gated K^+^ currents (N = 28, 2-way ANOVA compared to IGCs: p = 0.0001 main effect of cell type, p = 0.0059 interaction between cell type and stimulus intensity; 2-way ANOVA compared to EGCs: p = 0.052 main effect of cell type, p = 0.148 interaction between cell type and stimulus intensity). (**J**) Input-output curve for normalized voltage-gated K^+^ current inactivation (N = 28, 2-way ANOVA compared to IGCs: p = 0.0001 main effect of cell type, p = 0.001 interaction between cell type and stimulus intensity; 2-way ANOVA compared to EGCs: p = 0.054 main effect of cell type, p = 0.0752 interaction between cell type and stimulus intensity). (**K**) Spontaneous EPSC frequency (N = 27, p = 0.21 to IGCs p = 4.77 × 10^-6^ to EGCs by Wilcoxon rank sum test). (**N**) Spontaneous EPSC amplitude (N = 27, p = 2.61 × 10^-8^ to IGCs, p = 6.10 × 10^-4^ to EGCs by Wilcoxon rank sum test).

We observed two distinct morphological and electrophysiological types in the AOB GL, potentially including periglomerular cells (PGs) and some short axon cells (SAs, Fig. 6A-D). Some AOB JGCs had low capacitance and responded to moderate depolarization with sustained action potential generation (Fig. 6 B-C). These JGCs responded to depolarizing voltage steps with non-accommodating voltage-gated K^+^ currents, and often showed a slowly-activating K^+^ current (Fig. 6B-C). Other AOB PGs, by contrast, had higher capacitance and lower input resistance, suggesting a larger and/or more highly conductive dendritic arbor. These cells often responded to mild depolarization by generating plateau potentials (Fig. 6D). Plateau potentials in these cells were sometimes observed in the absence of somatic current injection, or following hyperpolarizing current injections, suggesting that spontaneous EPSPs and/or rebound depolarization were sufficient to drive these events. Unlike regular-spiking JGCs, plateau potential-generating JGCs responded to voltage-clamp depolarization with nearly-completely inactivating K^+^ currents (Fig. 6D). JGCs, taken as a whole, demonstrated mean physiological features that between IGCs and slightly lower than EGCs (Fig. 6E-L). This was likely a byproduct of the apparent split between putative PGs and SAs.

To evaluate the collective physiological differences between AOB JGCs, IGCs, and EGCs, and potentially associated with genetically labeled types, we introduced the physiological features of 26 JGCs and 9 LOTCs into the dataset containing IGCs, EGCs, and MCs (total 118 neurons) and re-clustered the data. This analysis identified 11 clusters and a spectrum of electrophysiological properties that spanned the diverse set of AOB interneurons (Fig. 7). As in previous clustering analyses, subdivisions within the JGC population became evident, with plateau potential-generating and regular-spiking JGCs associated with different clusters (Clusters 2 and 5, respectively; **Fig. A-C**). Cluster 1 from this analysis contained faster-spiking, but strongly accommodating cells lacking prominent I_H_ currents (4 EGCs, 1 IGC, and 2 JGCs) (Fig. 7D). Cluster 2 contained cells that spiked extremely weakly and showed rapid spike adaptation, and was enriched in plateau potential-generating JGCs (6/9 cells in cluster), all of which were labeled in *Calb2*-tdTomato transgenic lines (Fig. 7E). Clusters 3 and 4 were enriched in weakly-spiking IGCs (8/9 in Cluster 3; 13/17 in Cluster 4; Fig. 7F-G). Cluster 5 was enriched in regular-spiking JGCs (13/16 cells in cluster), all of which were labeled in *Gad2*-tdTomato transgenic mice (Fig. 7H). Thus, multidimensional analysis confirmed the apparent split of JGCs into regular-spiking and plateau potential-generating subgroups, and further revealed that the *Gad2*-tdTomato labeled JGCs were enriched in the regular-spiking phenotype while *Calb2*-tdTomato labeled JGCs were enriched in the plateau potential-generating phenotype. Clusters 6 and 7, similarly to Cluster 1, included a mix of faster-spiking IGCs and EGCs (Fig. 7I-J). Clusters 8 and 9 were enriched in EGCs of all transcriptional types (12/21 in Cluster 8; 6/9 in Cluster 9) but also included a substantial number of LOTCs (5/30 across the two clusters)(Fig. 7K-L). The apparent split among LOTCs into IGC/JGC-dominated clusters (4/9 LOTCs in Clusters 2-4) and EGC-dominated clusters (5/9 LOTCs in Clusters 8-9) indicates that LOTCs include a mix of IGC-like neurons and EGC-like neurons. As in other clustering runs, Clusters 10 and 11 were dominated by MCs (12/14 cells across the two clusters). Taken as a whole, clustering based on this set of intrinsic electrophysiological parameters provided useful information that identifies functional subdivisions within morphological classes and an objective mechanism for evaluating relationships between genetically-defined and morphologically-defined cell types.

**Figure 7.**
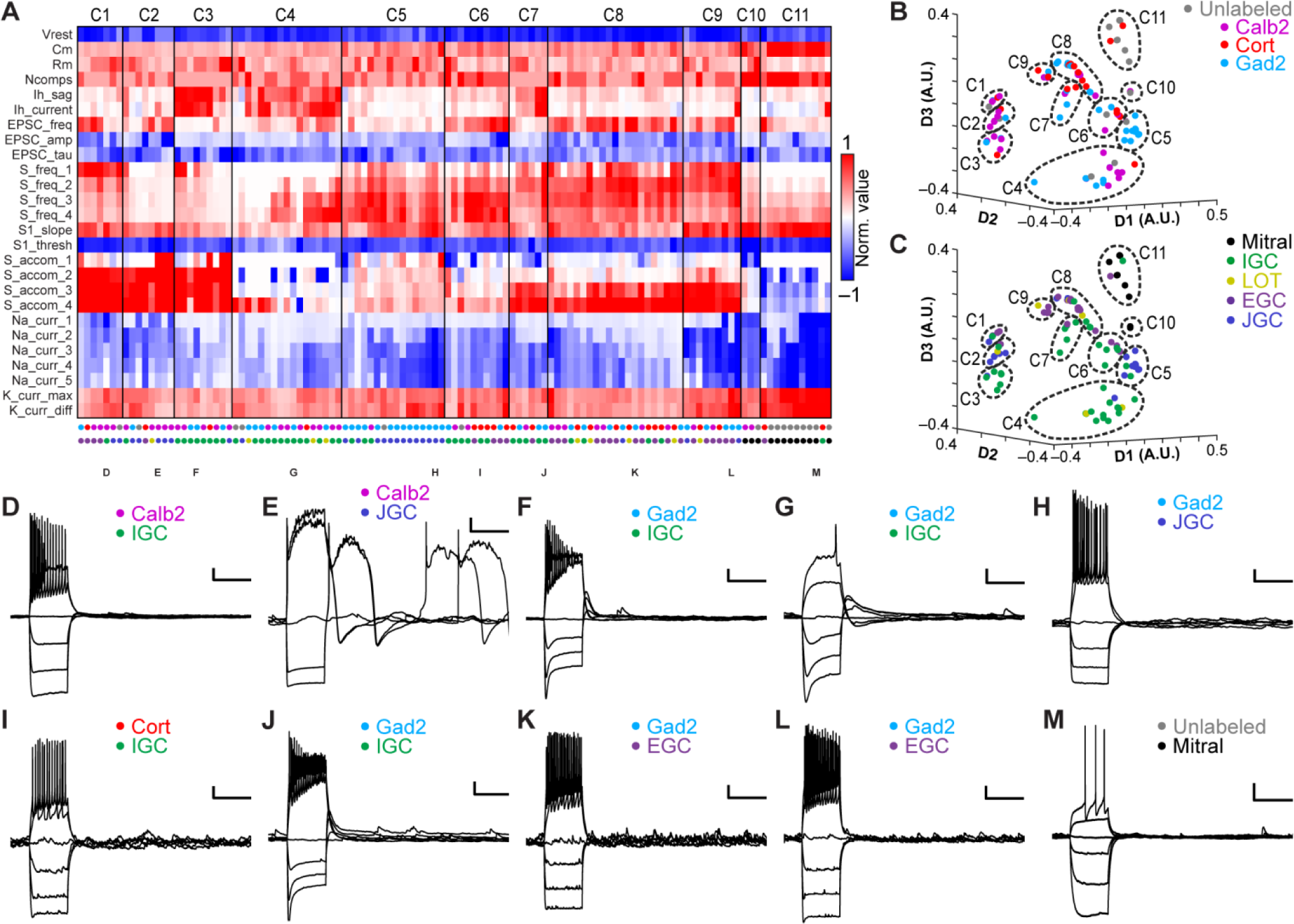
Multidimensional analysis across AOB neuronal types. (**A**) Cluster analysis of the 118 interneurons and mitral cells for which all 26 physiological properties were obtained. Normalized values were calculated as per Figure 3. Row labels refer to intrinsic properties listed in Table 1. Each column represents a single cell, and column labels are color coded based on genetically-defined type and morphologically-defined type. Clusters are separated by solid vertical lines. (**B-C**) Nonclassical multidimensional scaling of the cell properties shown in **A**, colorized by genetically-defined type (**B**) and morphologically-defined type (**C**). Dashed outlines indicate approximate cluster boundaries. C1-C11 refer to the cluster definitions in **A**. (**D-M**) Current clamp responses to the same current injection series shown in Figures 2, 4, and 6 from selected cells as noted below the columns in **A**. Scale bars: 10 mV, 500 ms.

## Discussion

### Transcriptional diversity among AOB interneurons is largely uncorrelated with intrinsic physiology

A major practical limitation to studying information processing in neural circuits is the available toolkit with which to selectively manipulate circuit elements. These tools can include electrical, optical, or pharmacological manipulations that selectively engage circuit elements at a particular point in space and/or time, and are increasingly involving conditional expression systems that leverage cell type-specific gene expression patterns. In a quest to improve upon our toolkit for studying the neural circuit logic of the AOB, we designed the studies presented here to be capable of objectively distinguishing between physiologically-distinct cell types within and across morphological and transcriptional categories. This general strategy is becoming increasingly important as novel technologies for combining electrophysiological and transcriptional data at the single cell level (Chen et al., 2016; Fuzik et al., 2016; Cadwell et al., 2017; Ting et al., 2018).

Taken as a whole, the results of these studies highlight the current challenges to circuit-level inquiries in understudied brain regions. Despite successfully identifying several physiologically distinct interneuron subpopulations in the AOB (especially within the morphologically-defined classes of IGCs and JGCs, Figs. 2, 3, 6, and 7), in only one case did we observe any specific segregation of genetic/transcriptional labels based on within-morphological-class physiological features (JGCs, Figs. 6–7). That said, the *Cort-cre* transgenic line, which labels interneurons sparsely throughout the brain, was almost fully selective for EGCs in the AOB (Fig. 1). The *Cort-cre* transgenic mouse line thus provides a compelling genetic tool for selective monitoring or manipulation of a distinct subtype of AOB interneurons; we expect this tool to be highly useful for future studies of AOB EGC function. Even so, the *Cort*-tdTomato EGC population is not pure (there are a few *Cort*-tdTomato IGCs and PGCs; Figs. 1–7), and it is not clear whether every AOB EGC is labeled in the *Cort*-cre transgenic line. Ultimately, these studies clearly indicate that new tools will be necessary for researchers to selectively monitor and manipulate functional groups of AOB interneurons. Such tools will almost certainly involve the identification of new marker genes, but may also involve the utilization of tools that selectively label interneurons based on recent activation (Cansler et al., 2017; Gao et al., 2017).

### IGC physiological diversity suggests multiple functionally-distinct subpopulations

IGCs are the most numerous and are currently the best-studied interneuron subtype in the AOB (Dudley and Moss, 1999; Jia et al., 1999; Araneda and Firestein, 2006; Smith et al., 2009; Taniguchi et al., 2012; Hu et al., 2016). Their known and hypothesized connections to experience-dependent plasticity make them a particularly compelling cell type to study (Brennan et al., 1990; Okutani et al., 1999; Cansler et al., 2017; Gao et al., 2017; Oboti et al., 2017). Despite the relatively strong set of inquiries into IGC function, there are many outstanding questions about IGCs’ role in AOB information processing and experience-dependent chemosensory plasticity. In this study, we did not focus on any specific aspect of IGC function, but instead explored the intrinsic physiological diversity within the IGC subtype in hopes of providing an improved quantitative foundation for future AOB studies.

The results of our systematic inquiry into IGC intrinsic features revealed at least 2, if not more, functionally-defined subpopulations of IGCs (Figs. 2–3). The most prominent differences between these subpopulations were the capacity to fire sustain action potential trains in the face of moderate depolarization and the magnitude of their HCN channel-mediated I_H_ sag potentials (Figs. 2–3). The functional implications of these results for AOB circuit function are not yet clear, but the data suggest – at a minimum – that AOB IGCs should not be considered to be a monolithic population. A substantial fraction of recorded AOB IGCs are poorly capable of generating more than one or two action potentials (Figs. 2D, 3). Given that AOB IGCs are replenished through adult-born neurons migrating through the rostral migratory stream (reviewed in Oboti and Peretto, 2014), it is possible that these IGCs – which are not selectively labeled by any of the 3 transgenic lines used in this study – represent immature (i.e. recently-arrived) adult-born neurons that have not yet fully incorporated into the AOB circuit. Even still, these IGCs exhibit spontaneous EPSCs of normal frequency and amplitude, suggesting they possess dendrites and functional synapses, and as such are theoretically capable of contributing to AOB processing even if they are “immature.” This feature – functional activity prior to full maturity – has also been seen in peripheral olfactory sensory neurons (Cheetham et al., 2016). Studying the conditions that bring about and contribute to neuronal maturation of IGCs seems likely to be an important component of inhibitory function in sensory processing (Oboti and Peretto, 2014).

Other functionally defined subpopulations of AOB IGCs include neurons that vary in their capacity to sustain high frequency spiking in the face of intermediate levels of somatic depolarization (Figs. 2–3). The expression of the prominent HCN channel-mediated I_H_ conductance in cells displaying intermediate intrinsic excitability (i.e. neurons capable of firing high frequency bursts, but readily entering depolarization block; Figs. 2C, 3) may indicate an intermediate functional phenotype among IGCs. This hypothesis is partially supported by recent studies that showed IGC intrinsic physiology is altered over 2-8 hour time courses by recent chemosensory-driven activity (Cansler et al., 2017; Gao et al., 2017). It is an intriguing possibility that these other, more subtle, differences in IGC physiology may reflect their involvement in recent activation. Given that AOB IGCs are hypothesized to contribute to experience-dependent chemosensory learning (Bruce and Parrott, 1960; Kaba and Keverne, 1988; Brennan et al., 1990; Okutani et al., 1999), it may be the case that the IGC physiological profiles established in this study are capable of distinguishing IGCs that span the range of immature, to mature but quiescent, to actively engaged IGCs.

### EGC physiological features suggest unique roles in AOB sensory processing

In the MOB and AOB, like most cortically-arranged brain circuits, interneurons populate major plexiform layers. In the MOB, parvalbumin-expressing interneurons in the external plexiform layer have been shown to possess unique physiological features that support divisive normalization (Kato et al., 2013; Miyamichi et al., 2013). Recent reports indicated a morphologically similar class of interneurons, EGCs, is present in the rodent AOB, and each of the transgenic lines utilized in this study labeled EGCs (Fig. 1). AOB EGCs possess broad, spinous dendrites that spread out along the anterior-posterior axis of the AOB (Larriva-Sahd, 2008)(Fig. 4). Based on morphology alone, one might hypothesize that this morphology would support the same type of broad innervation and gain-scaling function served by parvalbumin-expressing interneurons in the MOB. Since there are no previous reports of AOB EGC physiology, we were excited to explore the physiological features that, along with the morphological features of these cells (Fig. 4), might inform hypotheses about their function.

The intrinsic physiology of AOB EGCs was unique and in some ways extreme (Figs 4–5). Incredibly, AOB EGCs possessed resting membrane potentials, averaging around −0 mV, which were effectively at the Nernst potential for K^+^ ions with this combination of internal and bath solutions. These resting membrane potentials were the same as AOB astroglia (data not shown), and at the extreme end of expectations for neurons in the brain. The goal of this study was not necessarily to interrogate the specific physiological/biophysical mechanisms for this extreme phenotype, but did suggest that EGCs, at rest, are approximately 40 mV hyperpolarized to spike threshold, and may require extensive activation by MCs to fire action potentials. Intriguingly, another feature that distinguished AOB EGCs from other cells was their incredibly high rates and amplitudes of spontaneous glutamatergic EPSCs (Figs. 4, 5, and 7). This observation, coupled with the dense, spinous dendritic morphology of EGCs, indicates broad and dense innervation by MCs. AOB EGCs, unlike IGCs, possessed small I_H_ sag potentials, and readily sustained 20-40 Hz trains of action potentials upon moderate somatic depolarization (Figs. 4–5). This suggests that these cells are capable of distributing broad lateral inhibition among spatially-dispersed MCs, but further studies are necessary to investigate this hypothesis.

All three transgenic lines explored in this study labeled EGCs, which are dwarfed in total numbers by IGCs (Fig. 1). EGCs labeled in *Cort-cre* transgenic mice were indistinguishable from *Gad2-* and *Calb2-cre* labeled EGCs (Fig. 5), but were much less dense than those populations (Fig. 1). This suggests that EGCs labeled in *Cort-cre* transgenic mice represent a subset of the total EGC population. Cortistatin, a somatostatin-like neuropeptide hormone (de Lecea et al., 1996) has never been studied in olfaction; these studies indicate that its role in the AOB may be to modulate MC activity, but further studies are necessary to evaluate this hypothesis.

### AOB JGCs include physiologically-distinct subsets associated with marker gene expression

In the MOB, extensive studies into the role of JGCs have identified several JGC subtypes, including periglomerular cells (PGs) and short axon cells (SAs), which play important roles in inter-and intra-glomerular processes (McGann et al., 2005; Tan et al., 2010; Fukunaga et al., 2014; Livneh et al., 2014). PGs, in general, are thought to support intraglomerular processes, like presynaptic silencing of sensory input, and short-range interglomerular inhibition (reviewed in Burton, 2017). SAs, on the other hand, participate extensively in long-range interglomerular inhibition through their activation and inactivation of other cells, including MCs and external tufted cells. Thus, there is abundant evidence for functional diversification among JGCs in the MOB. In the AOB, however, there is very little available information about JGC function (Goldmakher and Moss, 2000; Hu et al., 2016).

The results of our survey of JGCs revealed the presence of two JGC subpopulations with macroscopically similar in their intrinsic physiological features to PG subsets in the MOB (Puopolo and Belluzzi, 1998; Murphy et al., 2005; Sethupathy et al., 2013; Najac et al., 2015; Burton et al., 2017)(Figs. 6–7). We found that *Gad2*-tdTomato positive cells were nearly all capable of firing sustained action potential trains upon moderate depolarization (Fig. 6B,C). These cells possessed low capacitance and high input resistance, and superficially resemble a faster-spiking populations of MOB PGCs (Puopolo and Belluzzi, 1998; Najac et al., 2015). *Calb2*-tdTomato JGCs demonstrated spontaneous and stimulated plateau potentials, similar to a major class of MOB PGCs (Murphy et al., 2005; Masurkar and Chen, 2011; Fogli Iseppe et al., 2016). We did not explicitly seek out cells with morphological features matching SAs, but we cannot rule out the possibility that a subpopulation of the regular-spiking *Gad2*-tdTomato JGCs were AOB SAs. In all, these data confirm the presence of physiologically-distinct classes of JGCs in the AOB GL. The AOB GL is macroscopically much different in its organization compared to the MOB. The AOB GL has numerous, small glomeruli per vomeronasal receptor input and lacks a clear periglomerular cell boundary surrounding individual glomeruli (Pinching and Powell, 1971; Belluscio et al., 1999; Rodriguez et al., 1999). As such, the specific computations provided by AOB JGCs are likely to be different than their MOB counterparts. That said, our results indicate that the AOB utilizes JGC subpopulations with similar intrinsic physiological features to their MOB counterparts, a finding which will support future inquiries into AOB JGC function.

### Implications of current work for models of AOB circuit function

We designed this set of experiments with the simple goal of producing a better quantitative foundation for studying the many populations of interneurons performing inhibition (and perhaps excitation) in the AOB. The results identify functional subdivisions within morphological types and highlight the need for improved tools for selective labeling of AOB interneurons. By subjecting each of the 150 cells included in these analyses to a simple panel of intrinsic physiological stimuli, we were able to use multidimensional analysis methods to identify patterns of features that were shared by groups of neurons. These stimulus panels and the physiological features extracted from the data are easy to implement (taking ~2 minutes of time after patching a cell), and can be included in any existing study in which internal patch solutions allow the observation of natural spiking patterns in current clamp (e.g. potassium gluconate-based solutions, etc.). Future studies of AOB interneurons *in vitro* or *in vivo* including these simple intrinsic “fingerprinting” protocols will be able to compare the physiological profiles observed in each cell to those observed in this study. This approach supports the identification of novel physiological types or specific changes to established functionally-defined types that may occur in response to chemosensory experience (Cansler et al., 2017) and/or neuromodulation (Smith et al., 2009; Smith and Araneda, 2010; Leszkowicz et al., 2012). Taken as a whole, these results provide a major improvement in the available physiological data for AOB interneurons, and improve the foundation for studies of AOB circuit function.

Author contributions
MAM, and JPM performed the electrophysiological recordings. JMT and MAM conducted immunohistochemical studies. JPM, HLC, KEZ, MAM, and DJR analyzed data. JPM wrote the manuscript.

## Acknowledgements

We thank Salma Ferdous, Natasha Browder, and Cara Nielson for technical support.

